# Modeling the Three-Dimensional Chromatin Structure from Hi-C Data with Transfer Learning

**DOI:** 10.1101/2021.12.15.472387

**Authors:** Tristan Meynier Georges, Maria Anna Rapsomaniki

## Abstract

Recent studies have revealed the importance of three-dimensional (3D) chromatin structure in the regulation of vital biological processes. Contrary to protein folding, no experimental procedure that can directly determine ground-truth 3D chromatin coordinates exists. Instead, chromatin conformation is studied implicitly using high-throughput chromosome conformation capture (Hi-C) methods that quantify the frequency of all pairwise chromatin contacts. Computational methods that infer the 3D chromatin structure from Hi-C data are thus unsupervised, and limited by the assumption that contact frequency determines Euclidean distance. Inspired by recent developments in deep learning, in this work we explore the idea of transfer learning to address the crucial lack of ground-truth data for 3D chromatin structure inference. We present a novel method, *Transfer learning Encoder for CHromatin 3D structure prediction (TECH-3D)* that combines transfer learning with creative data generation procedures to reconstruct chromatin structure. Our work outperforms previous deep learning attempts for chromatin structure inference and exhibits similar results as state-of-the-art algorithms on many tests, without making any assumptions on the relationship between contact frequencies and Euclidean distances. Above all, TECH-3D presents a highly creative and novel approach, paving the way for future deep learning models.

## 1 Introduction

Eukaryotic genomes are complex, long molecules that carry all hereditary information of an organism in their sequence. In human cells, the ∼2-meter-long DNA molecule resides in a nucleus of ∼10 *μ*m diameter [13]. To achieve this remarkable compression, DNA is packaged around proteins to form chromatin, which is in turn hierarchically organized in loops, topologically associated domains (TADs) and higher-order compartments [11]. This 3D organization allows the genome not only to fit within the confined nuclear space, but also to control *if, when* and *how* genetic information is accessed [2]. Indeed, a plethora of studies have highlighted that the 3D structure of chromatin is not random, but rather ensures timely and accurate completion of essential cellular processes, such as gene expression and DNA replication [18, 16]. Distortions of 3D chromatin structure have important implications in disease, leading to rewiring of regulatory interactions and linked to, among others, developmental defects, neurological disorders and, notably, a variety of cancer types [5, 10]. Hence, understanding 3D chromatin structure has the potential to not only lead to a mechanistic comprehension of biological processes, but also elucidate disease pathogenesis [1].

To study chromatin organization, early studies used imaging techniques such as fluorescence in situ hybridization (FISH) [4], which allows measurements of pairwise Euclidean distances between genomic loci. However, due to technical limitations in throughput and resolution, FISH-based methods cannot capture genome-wide organization. To fill this gap, the so-called high-throughput, high-resolution chromosome conformation capture technique (Hi-C) has been proposed [9]. In mcontrast to FISH, Hi-C does not measure Euclidean distances but *frequencies of physical interactions*. A typical Hi-C experiment quantifies interaction frequencies (IFs) between *N* pairs of tiny DNA fragments (high-resolution) across the entire genome (high-throughput), and can be summarized into an *N* ×*N* contact matrix (Figure 1). Hi-C can be performed either at a *single-cell* or *population* level. Since single-cell Hi-C contact matrices are sparse and noisy [19], inferring 3D chromatin structure from Hi-C data typically translates to recovering 3D structures from a single population Hi-C contact matrix (Figure 1).

**Figure 1:**
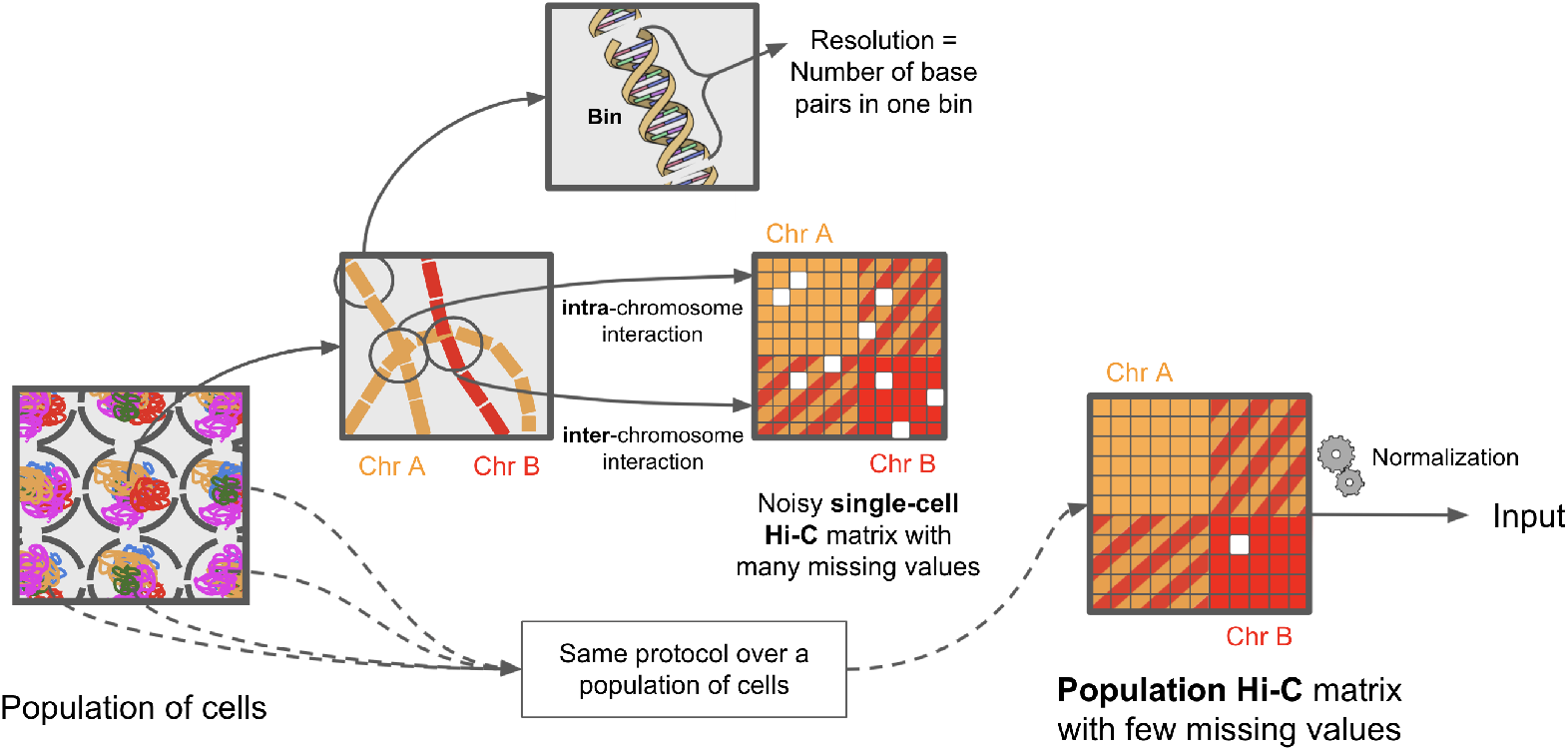
Schematic overview of Hi-C data generation.

### Related Work

Existing methods for 3D chromatin structure prediction take as input a population Hi-C matrix and either output a single *consensus* 3D structure, or an *ensemble* of possible 3D structures. Depending on how interaction frequencies are interpreted, these methods can also be categorized as *distance-, probability-* or *contact-based* [12]. Assuming that spatial distances and interaction frequencies are inversely correlated, distance- and probability-based methods first translate interactions to distances by employing a *transfer-function*. Then, the problem can be formulated either as traditional multidimensional scaling (MDS) [8] using the translated distances as constraints for distance-based methods (e.g., miniMDS [15]), or as probabilistic modeling for probability-based methods. On the other hand, contact-based methods directly map the interaction frequencies to an output structure(s) by employing manifold learning, with a notable example the method GEM [24] that exploits a modified version of stochastic neighborhood embedding (t-SNE) [22]. Recently, deep learning based methods have also been proposed. For example, REACH-3D [3] infers 3D chromatin structure by performing manifold learning on the Hi-C matrix using a recurrent neural network autoencoder.

However, current approaches suffer from several limitations. Firstly, with the exception of miniMDS that employs a divide and conquer strategy, existing methods are computationally demanding and do not scale well with high-resolution Hi-C data. Secondly, with the exception of GEM, most methods assume a fixed parametric transfer function mapping interaction frequencies to distances, but the exact mapping from interactions to distances differs between organisms and resolutions [23]. Thirdly, although REACH-3D circumvents both these limitations, its main drawback is the lack of ground-truth training data. Specifically, REACH-3D attempts to purposely overfit a single noisy Hi-C matrix, expecting that its latent representation can capture 3D structure. Since the model is unsupervised, its learning capacity is very limited and strongly depends on the noise level. Furthermore, REACH-3D ensures no control over the latent space, as its architecture cannot enforce it to actually encode for the 3D structure. The lack of ground truth data is thus a fundamental obstacle in the design of new deep learning models, not only limiting model validation and benchmarking, but also prohibiting the design of supervised approaches towards chromatin structure prediction. To address all these limitations, in this work we propose a novel deep learning approach called **Transfer learning Encoder for CHromatin 3D structure prediction (TECH-3D)**.

## 2 Methods

TECH-3D addresses the above limitations by exploiting *transfer learning*. We first go back to a more general and non-biological setting of **predicting *any* 3D structure from a synthetic contact frequency matrix**. This setting allows us to work in a (simpler) *supervised* setting and have complete control on the recovered structures. More formally, we define two problems:

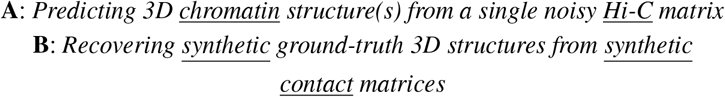

Although solving problem **A** is the goal of TECH-3D, problem **B** is easier to solve and, if the set of possible structures synthetically generated in **B** contains the set of true biological structures found in **A**, then **B** is more *generic* than **A**, a favorable condition to use transfer learning and solve **A** by exploiting **B**. Concretely, we develop a neural network Encoder as illustrated in Figure 2, trained on synthetic data from **B** and tested on experimental Hi-C matrices. The Encoder (Figure 2) first maps the synthetic Hi-C matrix to a *consensus* structure in the unit ball and then generates a corresponding distance matrix. We consider two types of neural network architectures for the Encoder: a Feed Forward network or a bidirectional Long Short-Term Memory (Bi-LSTM) network [17], as illustrated in the Appendix (Figures 8 and 9, respectively). Algorithm 1 summarizes the main steps of TECH-3D, and Sections 2.1 and 2.2 describe in detail the synthetic data generation and the loss function of TECH-3D, respectively.

**Figure 2:**
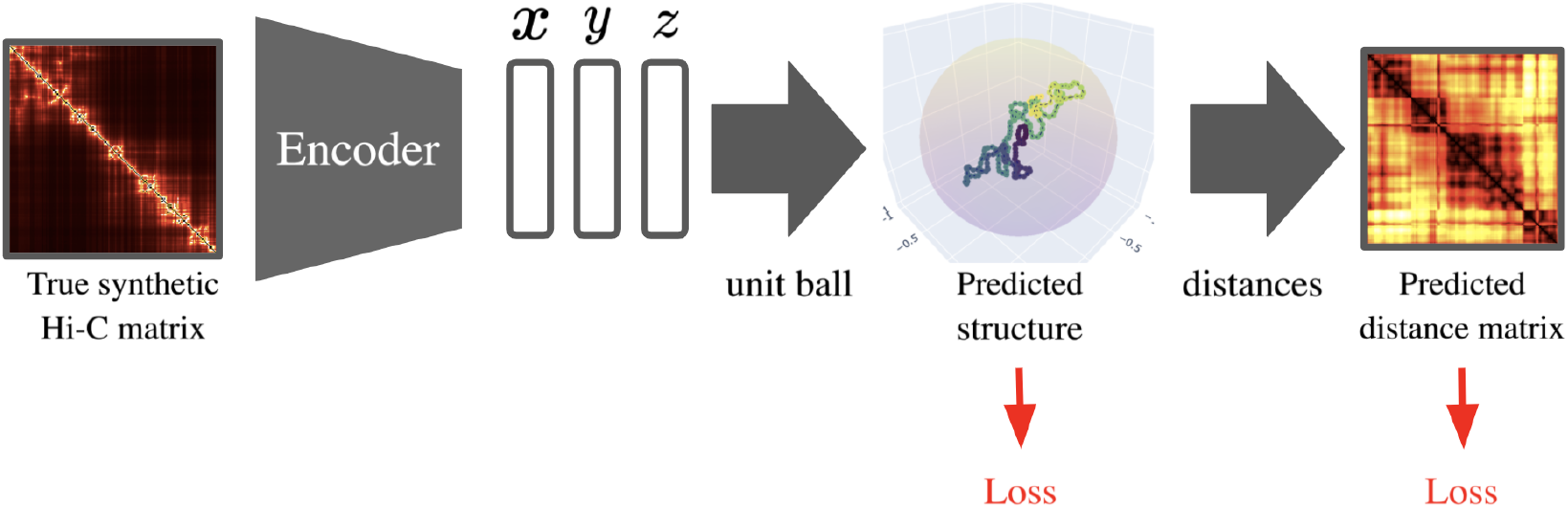
Overview of the TECH-3D implementation.

### Algorithm 1

TECH-3D Framework

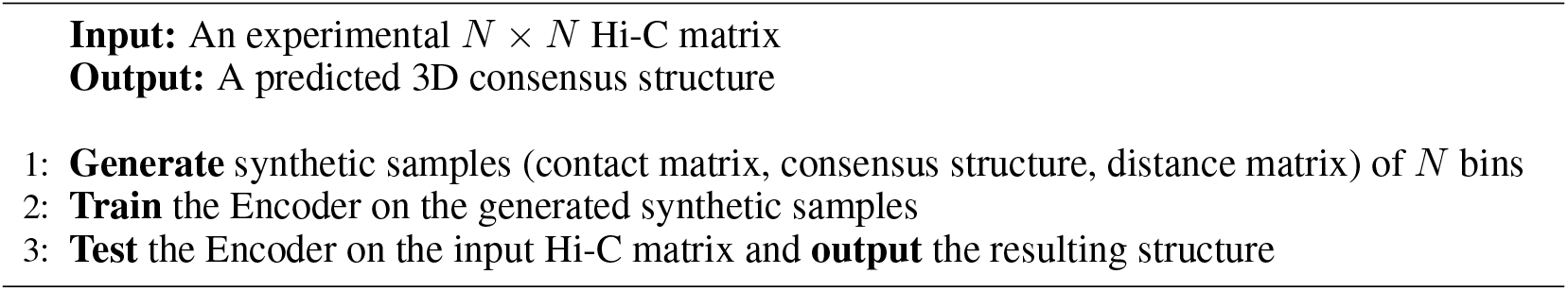

### 2.1 Synthetic Dataset

To our knowledge, the only publicly available synthetic dataset in the literature is the one developed by Trussart et al. [21] that contains 100 ground-truth 3D structures mapping to a single Hi-C matrix at various resolutions. The data is obtained by generating an ensemble of 100 hypothetical 1-megabase-long chromosome structures, from which a single synthetic Hi-C matrix is computed by mimicking the actual Hi-C protocol. Since in the Trussart data all 100 synthetic structures map to the same Hi-C matrix, we cannot exploit this dataset as training samples for problem **B**. For this reason, we designed our own procedure to generate synthetic data, where each structure comes with its own accompanying Hi-C matrix, as follows:

1: Produce a 3D structure *S*.

2: Compute the Euclidean distance matrix *D* from *S*.

3: Compute the contact matrix: 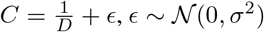.

We add noise to *C* such that the overall relationship between *C* and *D* resembles an inverse proportion while capturing biological variability. We consider two different heuristics to create structures, resulting in two different datasets, *B*_1_ and *B*_2_, described in detail in subsections 2.1.1 and 2.1.2, respectively. Since TECH-3D heavily depends on training data, we thoroughly investigated which training dataset performs the best when testing the model on problem **A**.

#### 2.1.1 Synthetic Random Walk Structures

We first considered Synthetic Random Walk Structures (*B*_1_), generated by uniformly sampling, at each step *N*, coordinates *x, y* and *z* from the set *S* of combinations [*i, j, k*], where *i, j, k* ∈ {−1, 0, 1}, with [0, 0, 0] excluded (i.e., *S* has size 3^3^ − 1). More formally:

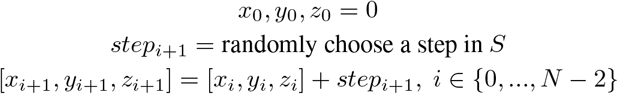

Figure 3 gives an example of a generated sample, that consists of pairs of synthetic 3D structures (a) with their corresponding Hi-C matrices (b). The Synthetic Random Walk Structure lacks loops and is less smooth than a real genome. Furthermore, the distance between consecutive bins is not constant (this should be approximately the case in a true Hi-C experiment). However, it still shares some common properties with genomes: it exhibits *in cis* interactions (diagonal of Figure 3b), cluster-like domains that resemble TADs (squares around the diagonal of Figure 3b) and long-range interactions (off-diagonal elements in Figure 3b), as well as an elongated overall shape.

**Figure 3:**
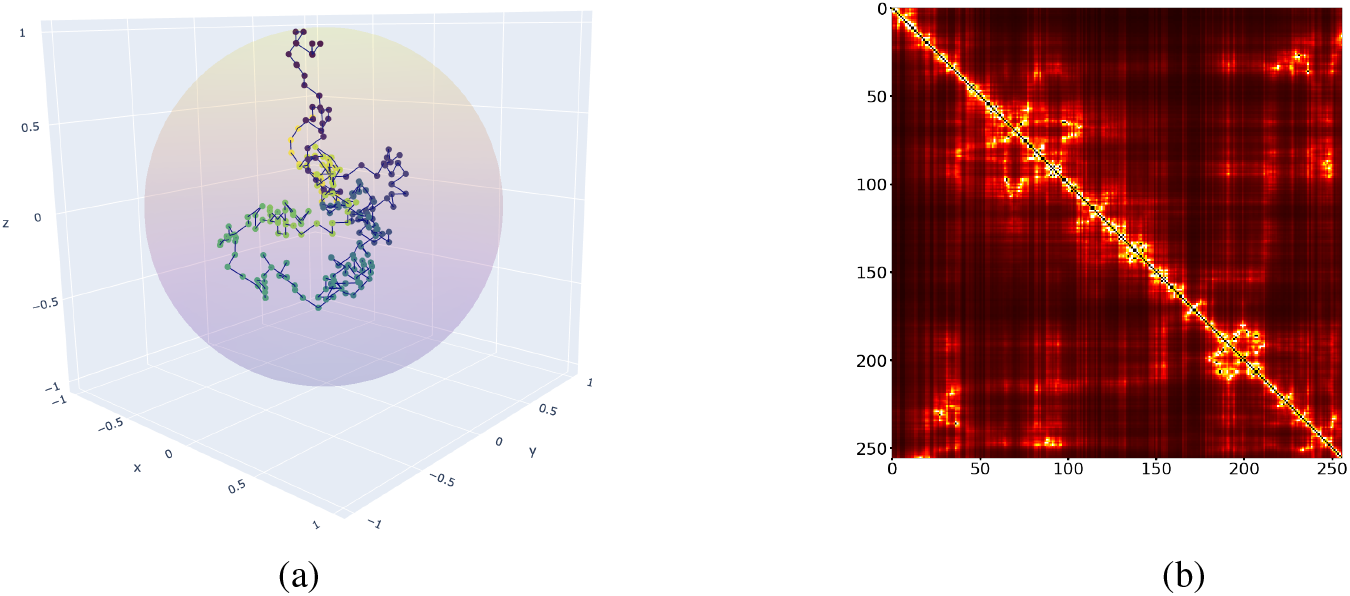
Output sample pair for the Synthetic Random Walk Structure heuristic for *N* = 256 bins.

#### 2.1.2 Synthetic Biological Structures

In this heuristic, we aim to generate 3D structures that are *biologically relevant* and more realistic than Synthetic Random Walks. In particular, we design Synthetic Biological Structures (*B*_2_) with a constant distance between consecutive bins and “smooth” loops. Starting from the origin [0, 0, 0], we generate coordinates as follows:

##### Algorithm 2

We denote by 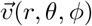 the vector starting from the origin and pointing to (*r, θ, ϕ*) in spherical coordinates (see Appendix Figure 10). Additionally, let 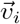 be the vector starting from bin *i* and pointing to bin *i* + 1 in the synthetic structure. To generate a structure, the algorithm recursively computes vectors 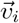 for *i* ∈ {0, …, *N* − 1} starting with 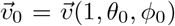, where the angles *θ*_0_ ∈ [0, *π*] and *ϕ*_0_ ∈ [0, 2*π*] are chosen uniformly. Now, 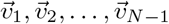 needs to be chosen so that conditions (1): constant distance between consecutive bins, and (2): “smooth” loops, are satisfied. Condition (1) translates to 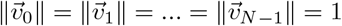. Satisfying condition (2) is more involved. If *_v*_*i*_ and *_v*_*j*_ are respectively the first and the last vectors of a loop, then:

- For *i* ≤ *k < j*, the angle 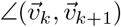 is small, resulting in smoothness inside the loop.
- For *i* ≤ *k < j*, the vector 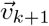 approximately points in the same direction as the previous vector 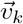 but with 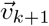 slightly more rotated towards the center of the loop than 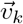 Formally, let *θ*_*k*_ and *ϕ*_*k*_ be the spherical coordinates of 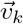 Then, 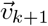 has coordinates *θ*_*k*_ + Δ*θ*_*k*+1_ and *ϕ*_*k*_ + Δ*ϕ*_*k*+1_, where *sign*(Δ*θ*_*i*_) = … = *sign*(Δ*θ*_*k*_) = *sign*(Δ*θ*_*k*+1_) = … = *sign*(Δ*θ*_*j*_), resulting in equal direction of rotation for all vectors in the loop. Similarly, *sign*(Δ*ϕ*_*i*_) = … = *sign*(Δ*ϕ*_*k*_) = *sign*(Δ*ϕ*_*k*+1_) = … = *sign*(Δ*ϕ*_*j*_).

When and how to create a new loop? We create a new loop by randomly changing the sign of Δ*θ*_*j*+1_ or Δ*ϕ*_*j*+1_ with small probability. By doing this, the vector 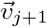 does not follow the same “loop pattern” as 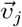 Concretely, the update rule is:

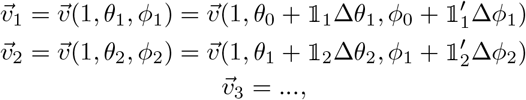

where: Δ*θ*_*i*_, Δ*ϕ*_*i*_ ∼ *U*_[0,*a*]_

and: 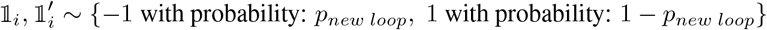

Parameters *p*_*new loop*_ and *a* control the compactness and smoothness of the generated structures, with larger values of *p*_*new loop*_ indicating stronger compactness and smaller values of *a* indicating higher smoothness. After experimenting with different values, we set 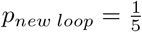 and *a* = 1.

One can notice that generating genome-like structures that preserve biological features (loops, constant distance between consecutive bins, etc.) quickly becomes complex. Figure 4 gives an example of an output sample. As desired, the Synthetic Biological Structure is smoother than the Random Walk Structure and exhibits loops, constant distance between consecutive bins, and an elongated overall shape. However, it does not contain clear TAD-like clusters, or long-range interactions.

**Figure 4:**
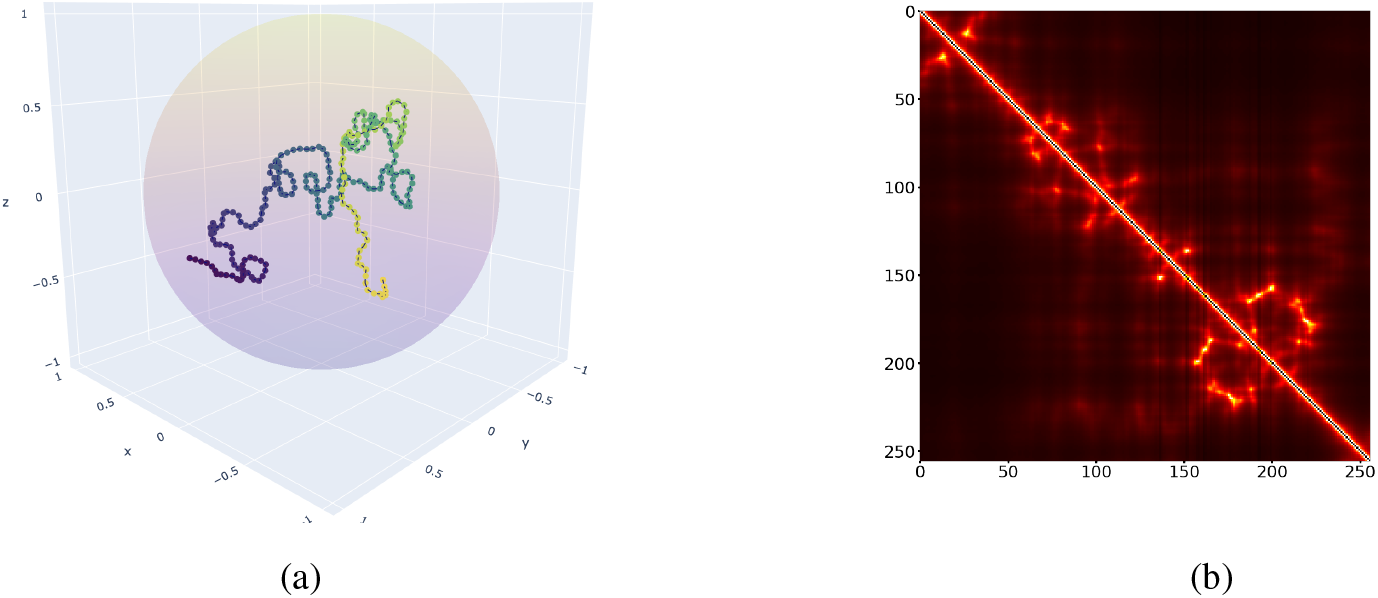
Output sample pair for the Synthetic Biological Structure heuristic for *N* = 256 bins.

### 2.2 Loss

The intuition for constructing an appropriate loss for TECH-3D is to measure the goodness of the predicted structure in terms of the distance between bins, the overall shape of the structure and its biological feasibility. The formula for the loss follows this idea and contains three terms for each feature, respectively:

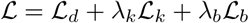

To test for the correctness of the distances between bins, we use the mean-squared error (MSE) between the true and predicted distance matrices *D* and 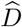:

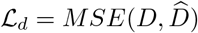

To test for the correctness of the overall shape, we compute the MSE between the true coordinates *S* and the predicted coordinates 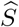, where the two structures are first aligned in space using the Kabsch algorithm [7]. This metric, called the Kabsch Distance, is a common metric to compare 3D structures in space:

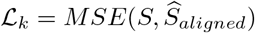

Finally, ℒ_*b*_ checks for the presence of biological features in the predicted structure, only relevant on Synthetic Biological Structures. Hence, by default *λ*_*b*_ = 0 when training on Synthetic Random Walk Structures. ℒ_*b*_ enforces the distance between consecutive bins to be approximately constant and the angle between consecutive vectors 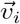 and 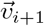 to be small for *i* ∈ {0, …, *N*−1} (where 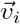 is the vector pointing from bin *i* to *i* + 1 in the sequence). The first objective is accomplished by minimizing 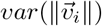 For the second, we penalize the dot product 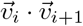 if it’ s negative. In this case, 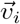 and 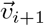 point in opposite directions, which induces a zigzagging, non-smooth structure.

Formally, we minimize 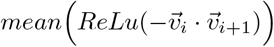 and the loss becomes:

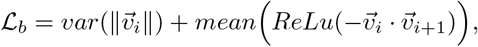

where 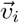 is computed from 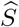.

## 3 Experiments and Results

### Datasets

The absence of real ground-truth biological structures necessitates alternative strategies for model training and validation. To circumvent this issue, we exploit our synthetic data generation framework with publicly available datasets for different tasks:

- **Our Synthetic Data:** The paired synthetic Hi-C matrices and structures generated in Section 2.1, used for TECH-3D training and optimization.
- **Trussart Synthetic Data:** The simulated dataset generated by Trussart et al. [21] (see Section 2.1), containing a single Hi-C matrix and 100 corresponding ground-truth structures (5 kilobase resolution), used for hyperparameter tuning and overall TECH-3D optimization. The Trussart structures are simulated so as to contain loops, TADs and long-range interactions, resembling real genomes.
- **Tanizawa Experimental Data:** The Fission Yeast dataset from Tanizawa et al. [20] that includes Hi-C measurements across the full fission yeast genome with a corresponding set of 18 microscopic FISH distances. We use the Hi-C matrix of 10 kilobases resolution and at timepoint *t*=20 minutes into the cell cycle for independent comparison with existing methods.

### Implementation details

The implementation is performed in PyTorch 1.9.0 [14]. As mentioned above, the network is first trained on our synthetic dataset that contains ground-truth structures, and the *weights* of the network are learned by gradient descent optimization. In particular, we use the PyTorch implementation of Adam [6] with the following parameters: a learning rate of 0.001, an exponential decay rate for the 1^st^ moment estimates of 0.9, an exponential decay rate for the 2^nd^ moment estimates of 0.999 and an epsilon value set to 10^*−*8^. Additionally, we train with early stopping for a number of epochs between 10 and 400. To implement early stopping, the base dataset is divided into 80% training and 20% testing samples. The dataset contains either 1000 samples (Sections 3.1 and 3.2) or 500 samples (Section 3.3) with a batch size of 10. The base dataset has no *validation* samples, since model optimization is performed on biologically relevant data, as explained in Section 3.1 below. The experiments were ran on a 2.7 GHz Intel Core i7 processor with 16 GB memory, in less than an hour. The source code is available here: https://github.com/PhosphorylatedRabbits/tech3d.

### 3.1 TECH-3D optimization

Given the many design choices of architecture, training dataset and hyperparameter values, we first need to optimize training of TECH-3D, so that it can generalize well to unseen, biologically relevant data. Specifically, we determine the hyperparameters *λ*_*k*_ and *λ*_*b*_ (Section 2.2) as well as the best combination of Encoder (Feed Forward or Bi-LSTM) and dataset (Synthetic Random Walk - *B*_1_ or Synthetic Biological - *B*_2_) through testing on the Trussart sample. Although, as discussed, the Trussart dataset cannot be used to train TECH-3D, it offers realistic structures that can be used for model selection and hyperparameter tuning while ensuring that TECH-3D generalizes well to real Hi-C data. The results for all combinations of architecture, dataset and *λ* values are summarized in Table 1, where the minimum test Kabsch Distance (KB) obtained on the Trussart dataset after training over 1 to 50 epochs is reported. We can observe that overall, the best architecture and dataset combination is a Feed-Forward Encoder trained on Synthetic Random Walk Structures, with *λ*_*k*_ = 0.1 showing the minimal Kabsch distance across all experiments. Bi-LSTM Encoders show an overall worse performance, potentially because they focus on a close neighborhood of the genome, which creates difficulties in predicting long-range interactions. For the same reason, Synthetic Random Walk Structures constitute the best training dataset, as they contain more long-range interactions. Based on these results, for the remaining of this section we fix TECH-3D as a Feed Forward Encoder and the training set as Synthetic Random Walk Structures.

**Table 1:** Min Kabsch Distance (KB) obtained on the Trussart test sample at 5 kilobase resolution for various hyperpameter values, architectures and train dataset types.

|  |  | Architecture |  |  |  |  |  |
| --- | --- | --- | --- | --- | --- | --- | --- |
|  |  | Feed Forward |  |  | Bi-LSTM |  |  |
| | | $\lambda_k$ | $\lambda_b$ | Min KB | $\lambda_k$ | $\lambda_b$ | Min KB |
| Synthetic Training Datasets | Random Walk | 0 | 0 | 0.0699 | 0 | 0 | 0.0870 |
|  |  | 0.1 | 0 | **0.0660** | 0.1 | 0 | 0.0883 |
|  |  | 1 | 0 | 0.0768 | 1 | 0 | 0.1227 |
|  |  | 10 | 0 | 0.1583 | 10 | 0 | 0.1707 |
|  | Biological | 0 | 0 | 0.0837 | 0 | 0 | 0.2053 |
|  |  | 1 | 0 | 0.0974 | 1 | 0 | 0.2325 |
|  |  | 1 | 1 | 0.0920 | 1 | 1 | 0.2316 |
|  |  | 10 | 1 | 0.1583 | 10 | 1 | 0.1729 |
|  |  | 10 | 10 | 0.1729 | 10 | 10 | 0.1757 |

### 3.2 Visualization of Predicted Structures

#### Predictions on our synthetic data

After model optimization, we first analyse the structures predicted on our synthetic data. Although the performance on our synthetic data is not relevant from a biological point of view, it provides insights on the complexity of the problem. As seen in Figure 5, the 3D structure predicted by TECH-3D (Figure 5b) highly resembles the the original synthetic structure (Figure 5a). Whereas details are harder to predict because the loss becomes too small, the distance matrix is almost perfectly reconstructed (Figures 5c - 5e). The Kabsch distance on our synthetic structures is on average 6 times smaller than on the Trussart sample (reported in Table 1).

**Figure 5:**
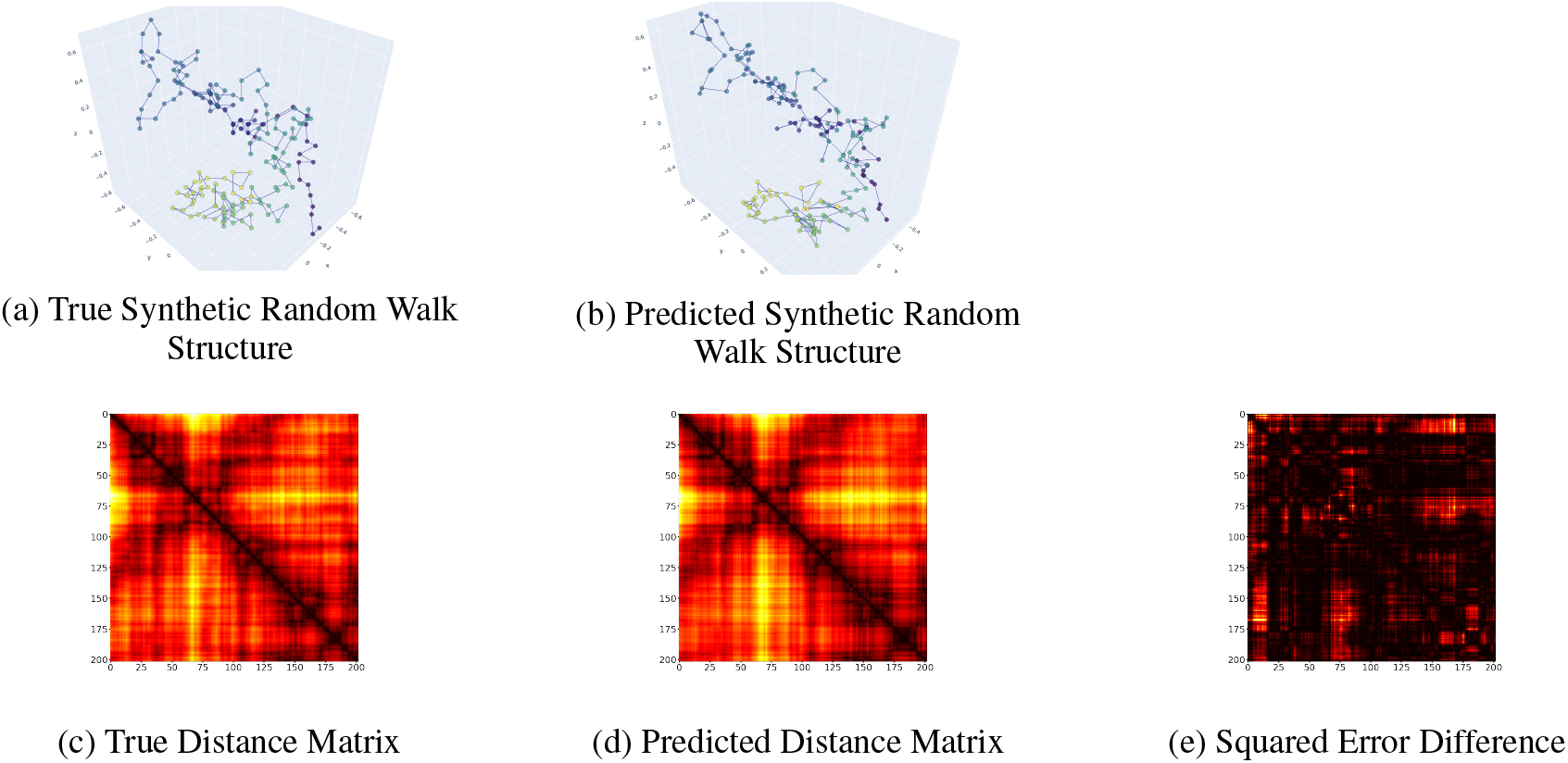
Ground-truth Synthetic Random Walk sample (left), TECH-3D prediction (middle) and squared error difference between the true and predicted distance matrices (right).

#### Predictions in the Trussart synthetic data

We then assess the performance of TECH-3D on the Trussart synthetic data. Since the 100 Trussart structures are highly similar, we compute their mean (see *Mean Trussart Consensus* in Figure 6) to simplify visualizations. Figure 6 displays the ground-truth structure (upper-left), the TECH-3D prediction, the output of several benchmark algorithms (namely REACH-3D, GEM and miniMDS) and random predictions. TECH-3D shows the best performance in terms of Kabsch distance, followed by REACH-3D, GEM and miniMDS. Furthermore, TECH-3D accurately predicts the overall structure, particularly within TADs where complex patterns are reconstructed with detail. However, the predicted structure is not smooth and the distance between consecutive bins is not approximately constant. Surprisingly, miniMDS captures both those features, but its Kabsch distance is high since the predictions within the TADs are noticeably wrong. This is because the Kabsch distance tends to favor predictions where TADs are simply clusters of points, as it is the case for GEM and REACH-3D.

**Figure 6:**
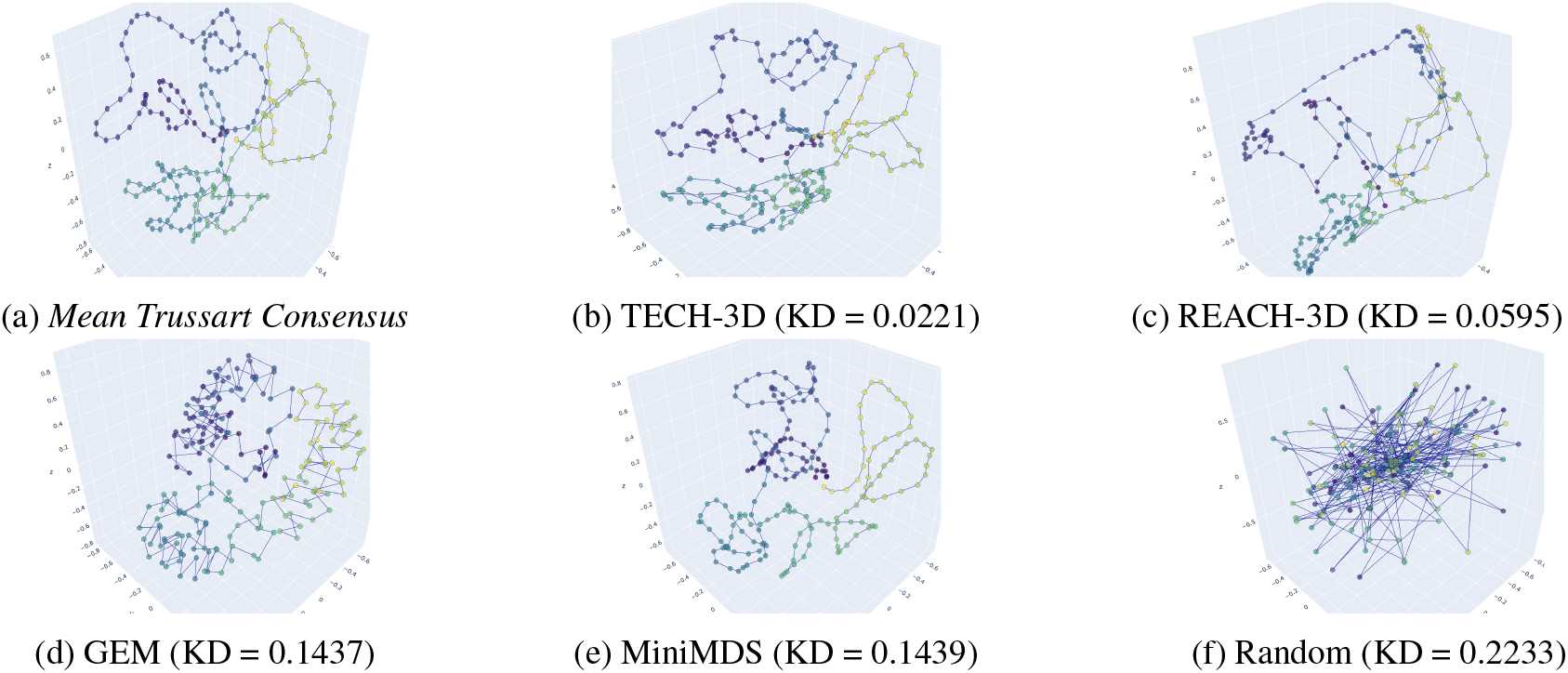
Output of various algorithms, ordered by Kabsch Distance (KD) with respect to the *Mean Trussart Consensus*.

Last, Figure 7 illustrates the distance matrices obtained from two different deep learning approaches: REACH-3D (unsupervised manifold-learning method with recurrent autoencoder) and TECH-3D (supervised method with transfer learning). We observe that REACH-3D clearly clusters the points within the TADs, as illustrated by the black squares on the distance matrix (Figure 7b). This is characteristic of unsupervised learning, where bins are simply clusters of points inside the 3D latent space to minimize the reconstruction loss. As we expected, transfer learning (TECH-3D) clearly outperforms unsupervised learning (REACH-3D).

**Figure 7:**
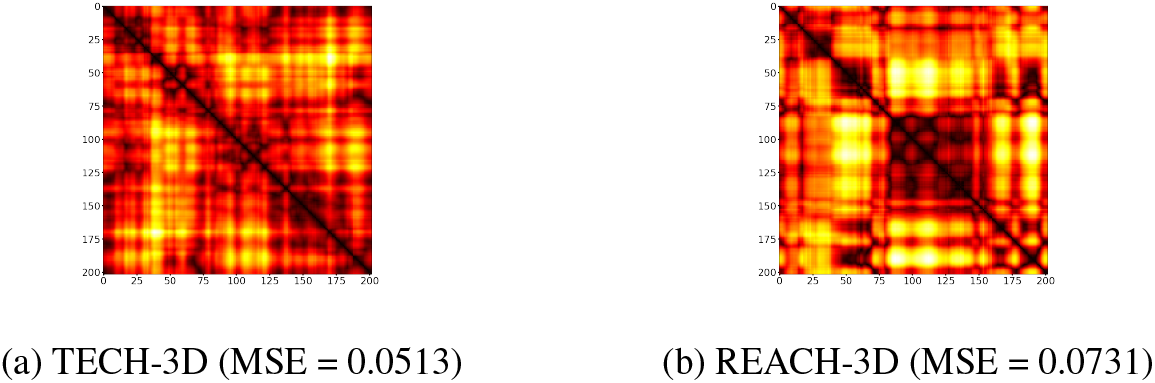
TECH-3D and REACH-3D output distance matrices. MSE with respect to the ground-truth *Mean Trussart Consensus* distance matrix given in the sublegends.

### 3.3 Evaluation on Experimental Data

Finally, we evaluate the TECH-3D performance on true experimental Hi-C data on the Tanizawa dataset. Since here no ground-truth structure is available, we use a small set of 18 genome-wide FISH distances experimentally measured by microscopy. Although the number of distances is very limited, by computing the correlation between actual FISH distances and predicted distances one can access the consistency of the prediction in a real genome. As seen in Table 2, both state-of-the-art algorithms GEM and miniMDS show very high correlation scores. The recent deep learning solutions have lower scores, but we observe a huge improvement from REACH-3D to TECH-3D — illustrating the higher learning capacity of transfer learning versus unsupervised learning.

**Table 2:**
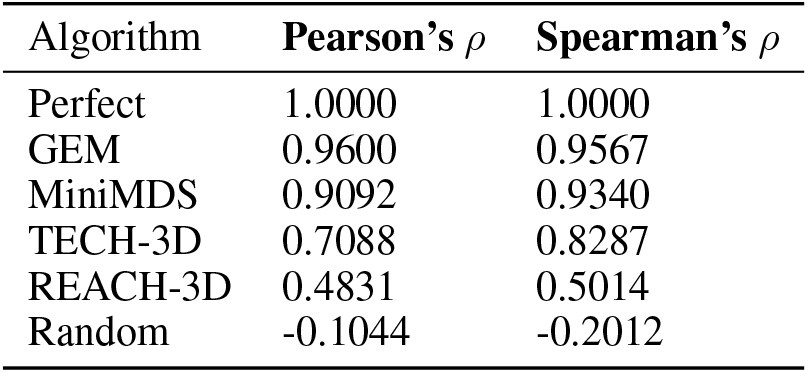
Correlation coefficients *ρ* between FISH and predicted distances for various algorithms

| Algorithm | Pearson’s $\rho$ | Spearman’s $\rho$ |
| --- | --- | --- |
| Perfect | 1.0000 | 1.0000 |
| GEM | 0.9600 | 0.9567 |
| MiniMDS | 0.9092 | 0.9340 |
| TECH-3D | 0.7088 | 0.8287 |
| REACH-3D | 0.4831 | 0.5014 |
| Random | -0.1044 | -0.2012 |

## Conclusions

In this work, we describe a creative and novel deep learning approach for genome structure inference, Transfer learning Encoder for CHromatin 3D structure prediction (TECH-3D). Our method employs transfer learning to overcome the lack of ground-truth data in the HiC setting — a major obstacle to the use of machine learning for this problem. When tested on publicly available synthetic structures, TECH-3D can accurately reconstruct them, outperforming state-of-the-art methods. When tested on microscopy measurements from real genomes, TECH-3D shows comparable performance as state-of-the-art algorithms and outperforms previous deep learning models. Above all, our work aims to present a highly creative perspective to the problem of inferring realistic 3D genome structures from Hi-C data, paving the way for future deep learning research in the field. Despite the methodological advancements achieved by TECH-3D, some aspects of the framework can be subject to improvement. Data generation is computationally intensive and can be enhanced by taking a divide-and-conquer approach, similar to miniMDS. Another promising direction could be to investigate new datasets, especially recent single cell Hi-C data (e.g., Dip-C [19], providing contacts for both sets of chromosomes (diploid) at a single-cell level.

## A Appendix

**Figure 8:**
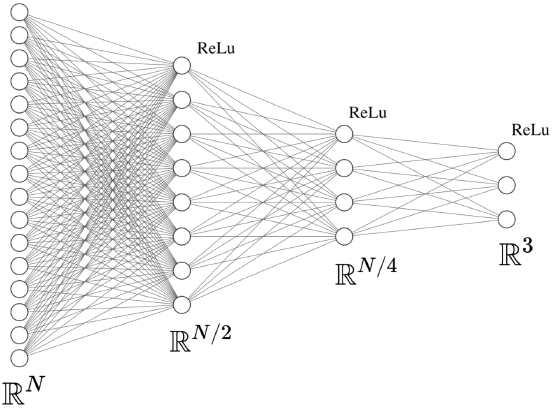
The Feed Forward Encoder architecture (example for *N* = 16). It applies three linear layers and a *ReLu* non-linearity function after each of them.

**Figure 9:**
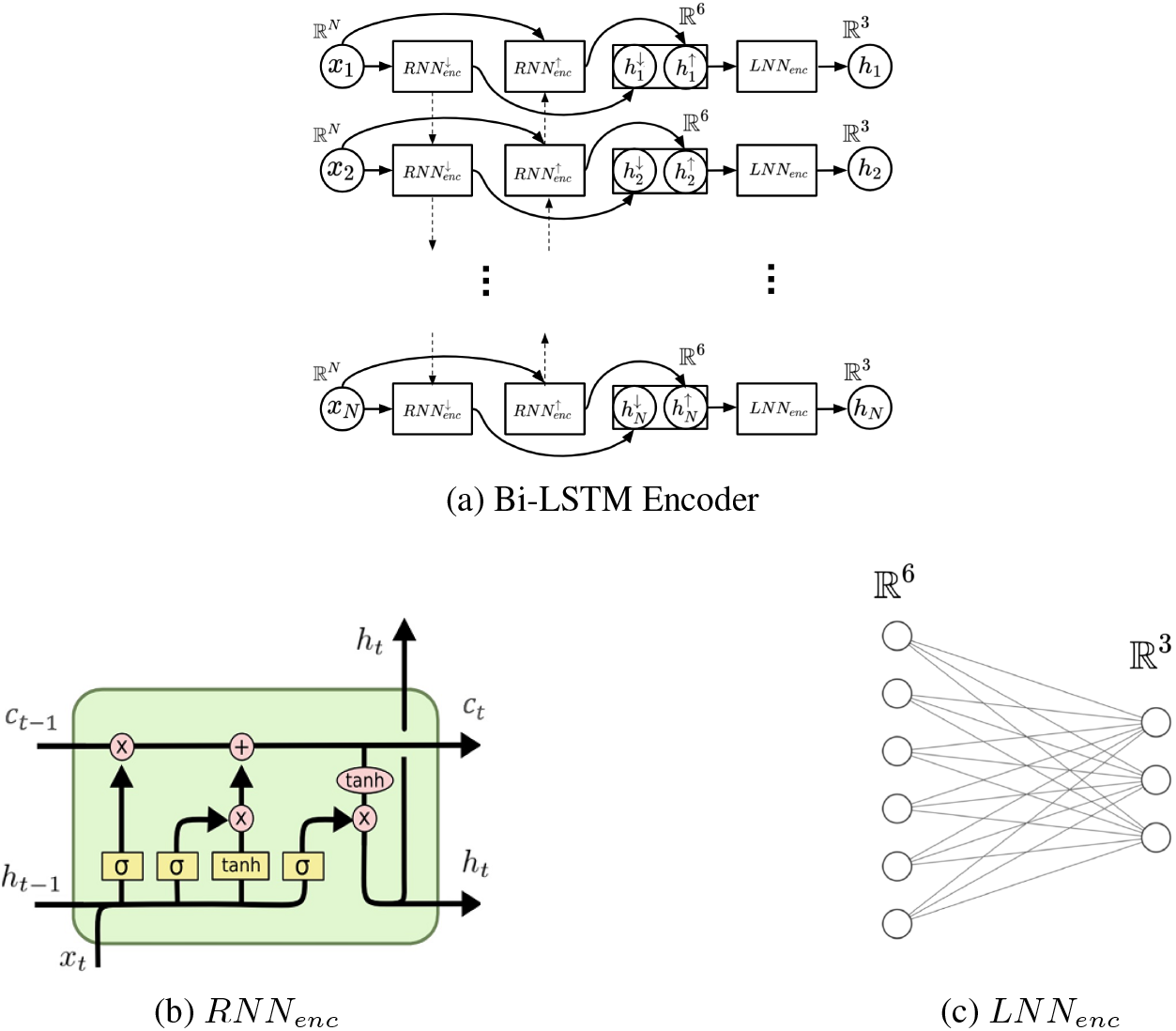
Overview of the Bi-LSTM Encoder architecture where *RNN*_*enc*_ is a LSTM cell (Figure 9b taken from Christofer Olah’ s blog (source: https://colah.github.io/posts/2015-08-Understanding-LSTMs/) and *LNN*_*enc*_ a linear network.

**Figure 10:**
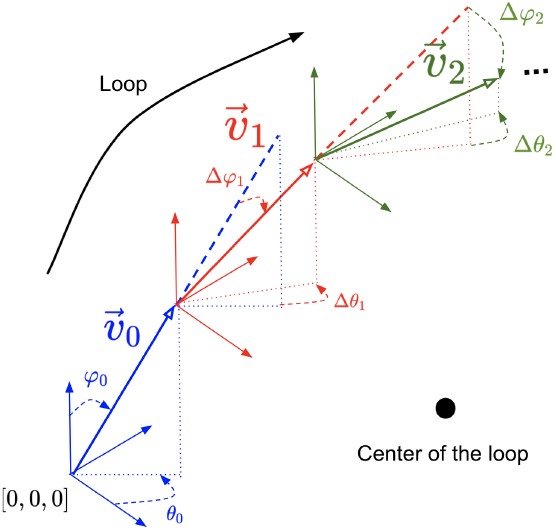
Visual example of the first steps of a *Synthetic Biological Structure* generation.

